# Acquisition of visual priors and induced hallucinations in chronic schizophrenia

**DOI:** 10.1101/498568

**Authors:** Vincent Valton, Povilas Karvelis, Katie L. Richards, Aaron R. Seitz, Stephen M. Lawrie, Peggy Seriès

## Abstract

Prominent theories suggest that symptoms of schizophrenia stem from learning deficiencies resulting in distorted internal models of the world. To further test these theories, we here use a visual statistical learning task known to induce rapid implicit learning of the stimulus statistics (Chalk *et al*., 2010). In this task, participants are presented with a field of coherently moving dots and need to report the presented direction of the dots (estimation task) and whether they saw any dots or not (detection task). Two of the directions were more frequently presented than the others. In controls, the implicit acquisition of the stimuli statistics influences their perception in two ways: 1-motion directions are perceived as being more similar to the most frequently presented directions than they really are (estimation biases); 2-in the absence of stimuli, participants sometimes report perceiving the most frequently presented directions (a form of hallucinations). Such behaviour is consistent with probabilistic inference, i.e. combining learnt perceptual priors with sensory evidence. We investigated whether patients with chronic, stable, treated schizophrenia (n=20) differ from controls (n=23) in the acquisition of the perceptual priors and/or their influence on perception. We found that, although patients were slower than controls, they showed comparable acquisition of perceptual priors, correctly approximating the stimulus statistics. This suggests that patients have no statistical learning deficits in our task. This may reflect our patients relative wellbeing on antipsychotic medication. Intriguingly, however, patients made significantly fewer hallucinations of the most frequently presented directions than controls and fewer prior-based lapse estimations. This suggests that prior expectations had less influence on patients’ perception than on controls when stimuli were absent or below perceptual threshold.

## Introduction

An increasingly popular idea in neuroscience is that perception and decision-making can be well described in terms of probabilistic inference processes (Knill and Pouget, 2004; Fiser *et al*., 2010; Friston, 2010; 2012). For example, statistical and perceptual learning studies show that the perceptual systems continuously extract and learn statistical regularities of the environment (for a review in visual perception, see e.g. Seriès and Seitz, 2013). This learning results in the construction of internal models of the environment, or expectations, which are used automatically and unconsciously to predict and disambiguate perceptual inputs in situations of uncertainty and to guide decisions.

In this context, it has been proposed that psychiatric disorders in general, and schizophrenia in particular, might be explained in terms of deficits in probabilistic inference (Friston, 2005; Frith and Friston, 2012; Adams *et al*., 2013; Jardri and Denève, 2013, for reviews see: Friston *et al*. 2016; Valton *et al*., 2017; Sterzer et al 2018). Impaired statistical learning and/or inference deficits would lead to distorted internal models of the world, which could then explain the existence of abnormal beliefs or delusions experienced by patients with schizophrenia. Incorrect perceptual inference may also lead to severe forms of illusions and result in the complex hallucinations that are part of the positive symptoms of schizophrenia.

Two different lines of research support this general idea. First, a number of studies using explicit probabilistic learning tasks, such as the “beads task” report that patients with schizophrenia show a deficit in integrating probabilistic information resulting in faster responses than control subjects, an effect called the ‘jumping-to-conclusions’ bias (Huq *et al*., 1988; Speechley *et al*., 2010; Averbeck *et al*., 2011; Evans *et al*., 2012). Interestingly, patients with stronger delusional symptoms fare worse at the task than those who do not (Huq *et al*., 1988; Speechley *et al*., 2010), and control subjects displaying delusional ideation also show similar impairments at the task (Freeman *et al*., 2008), suggesting a link between delusions and probabilistic inference (Garety *et al*., 2013; Garety and Freeman 2013). Second, patients with schizophrenia do not experience visual illusions in the same way as controls do. For example, patients are less susceptible to certain visual illusions such as the hollow-mask illusion (Dima *et al*., 2009; 2010; Keane *et al*., 2013; for review see: Silverstein and Keane 2011a; 2011b; Notredame *et al*., 2014). This suggests that they either have different implicit expectations about the environment (i.e. they would not have such a strong expectation that faces are convex) or that these expectations do not affect patients’ perception in the same way as observed in controls.

A few studies have recently tried to test the impaired Bayesian inference hypothesis more directly, but the findings are mixed. Teufel et al. (2015) found that early psychosis and schizotypal traits were associated with an increased influence of prior knowledge when disambiguating two-tone images. Powers et al. (2017) also reported an increased perceptual prior influence in experimentally-induced hallucinations in both patients and controls with higher propensity to hallucinatory experiences. Interestingly, however, a series of studies by Schmack et al. (2013; 2015; 2017) found an increased influence of cognitive priors on the perception of bi-stable stimuli in participants with schizotypal traits and clinical schizophrenia, but showed on the contrary a decreased influence of perceptual priors, suggesting that the level at which the prior operates in the inference hierarchy might lead to differential effects. Finally, Jardri et al. (2017) investigated probabilistic reasoning and found schizophrenia patients to be over-counting sensory evidence and under-weighting priors, which they described using a circular inference model. Together these findings paint a complicated picture of Bayesian inference in schizophrenia where priors can have either increased or decreased influence depending on the task, the stimulus and the type of priors involved (e.g. low-level perceptual prior *vs*. high-level cognitive prior).

A general limitation of these studies however, is that it is typically unclear to what extent the effects are driven by deficits in statistical learning (i.e. forming and updating the priors) or impaired inference *per se*. Moreover, past studies were usually only qualitatively comparing the behavioural results they collected with the proposed Bayesian theories.

To address these issues, we simultaneously investigated the implicit acquisition of priors, how these priors are integrated with sensory information and the influence they have on perception when stimulus is absent (i.e. experimentally induced hallucinations) in patients with schizophrenia. We used a previously developed statistical learning task (Chalk *et al*., 2010; Gekas *et al*., 2013; Karvelis *et al*., 2018) that is known to induce the rapid acquisition of the statistics of motion stimuli. In this task, participants need to report the direction of motion of a cloud of dots (estimation task) and whether they have perceived the dots or not (detection task; on some trials no stimulus is presented). Unbeknownst to the participants, two directions of motion are more frequently presented than others. Participants implicitly and unconsciously learn those stimulus statistics. This learning influences perception such that: 1) motion stimuli are perceived as being more similar to the most frequently presented stimuli than they really are (i.e. estimation biases); 2) participants sometimes report perceiving the most frequently presented stimuli in absence of visual stimuli (a form of hallucination). In previous work (Chalk *et al* (2010) Karvelis et al (2018)), we showed that Bayesian modelling could be applied to individual participants’ performances to quantitatively monitor their acquisition and use of the statistics of the stimuli (perceptual prior). We apply the same techniques in the current study to compare the perceptual priors acquired by patients with schizophrenia to those of controls.

## Materials & Methods

### Participants

A sample of 25 (22 male) individuals with psychosis (diagnosed with either DSM-IV schizophrenia, *n* = 21; or schizoaffective disorder, *n* = 4), and 23 (13 male) controls with normal or corrected-to-normal vision were recruited. Patients were recruited from inpatient and outpatient adult mental health services across NHS Lothian. Diagnoses were determined using the Structured Clinical Interview for DSM-IV (SCID-I; First, Gibbons, Spitzer, & Williams, 2002). None of the control participants met DSM-IV criteria for a psychotic disorder, bipolar disorder, or schizotypal or schizoid personality disorder. Symptom severity was measured with the Positive and Negative Syndrome Scale (PANSS) (Kay, Fiszbein, & Opler, 1987), current IQ with the Wechsler Abbreviated Scale of Intelligence (WASI; Wechsler, 1999) and pre-morbid IQ with the National Adult Reading Test (NART; Nelson, & Willison, 1991). All patients were medicated (85% on second generation anti-psychotics, 50% of these were also on mood stabilisers). The study was conducted in accordance with the national and international ethical standards for human experimentation and research (Declaration of Helsinki, 2013; Good Clinical Practice, 2014). All participants provided fully informed written consent. The study received ethical approval from the South East Scotland Research Ethics Committee 01 and NHS Lothian Research & Development. Following previous studies using the same paradigm (Chalk *et al*. 2010), we determined that 20 participants per group would give us ≥80% power to detect the correct acquisition of the prior and significance between groups (see power calculations in supplemental material). We thus aimed to recruit between 20-25 participant per group to account for possible exclusions due to poor performance at the task.

Twenty patients and twenty three controls successfully performed the task (see below and Supplementary Figure 1 for exclusion criteria based on performance). The demographic details of the included participants are shown in Table 1. It is of note that the patients all had relatively low levels of symptoms on the PANSS, but ongoing functional impairment.

**Table 1.** Participants’ demographics. PANSS = Positive and Negative Symptom Scale (lower score is better), GAF = Global Assessment of Functioning (higher score is better), OLZ eq. = Olanzapine equivalent dosage in mg/day. Values indicate mean and (standard deviation). For gender, group comparisons were done using Chi-square test, for all other measures, two-sided Wilcoxon rank-sum test was used.

| Characteristics | Controls (n=23) | Patients with schizophrenia (n=20) | Significance level p-value |
| --- | --- | --- | --- |
| Gender (males) | 13 | 17 | 0.04 |
| Age (years) | 33.86 (12.08) | 39.40 (9.43) | 0.07 |
| Premorbid IQ | 115.55 (4.41) | 113.18 (8.76) | 0.58 |
| Current IQ | 117.45 (7.28) | 111.15 (10.70) | 0.06 |
| PANSS Positive Scale | 8.86 (2.29) | 12.55 (5.01) | <0.01 |
| PANSS Negative Scale | 8.41 (2.24) | 12.20 (5.05) | <0.01 |
| PANSS General Scale | 22.27 (6.48) | 26.10 (8.35) | 0.12 |
| PANSS Total | 39.55 (9.22) | 50.85 (15.86) | <0.01 |
| GAF | 74.81 (11.42) | 54.75 (14.36) | <0.001 |
| OLZ eq. (mg/day) | --- | 12.61 (6.24) | N/A |
| Illness duration (years) | --- | 13.33 (9.01) | N/A |

### Apparatus, Stimuli & Procedure

The setup for this study was similar to that used by Chalk *et al*. (2010). Motion stimuli consisted of a field of dots with a density of 2 dots/deg^2^, moving coherently (100%) at a speed of 9°/s.

Each trial was composed of two tasks arranged as follows (**Fig. 1A**): First, participants were presented with a fixation point (0.5° diameter) for 400 ms. With the fixation point still on-screen, the motion stimulus (field of dots) was displayed along with a red bar extending from this fixation point. During the presentation of the field of dots, participants were required to estimate the direction of motion by aligning the red bar into the perceived direction of motion (*Estimation task*). The angle of this bar was randomized on each trial and participants were instructed to focus their gaze on the fixation point throughout the estimation task. The display then cleared when either the participant clicked the mouse to validate their choice (estimation) or 3000 ms had elapsed. After the estimation, a 200 ms delay was enforced before the detection screen was presented. The new screen was divided in two equal areas reading ‘Dots’ and ‘No Dots’, giving the participants a two-alternative forced choice (2-AFC). Participants were required to move the cursor to the right or the left to indicate whether they detected dots or not, and click to validate their choice (*Detection task*). The cursor then flashed green or red for correct or incorrect responses respectively. No time-outs were enforced during the detection task. Finally, the screen was cleared for 400 ms before a new trial began. Every 20 trials, participants were presented with feedback on their estimation performance in terms of average estimation error (in degrees).

**Figure 1:**
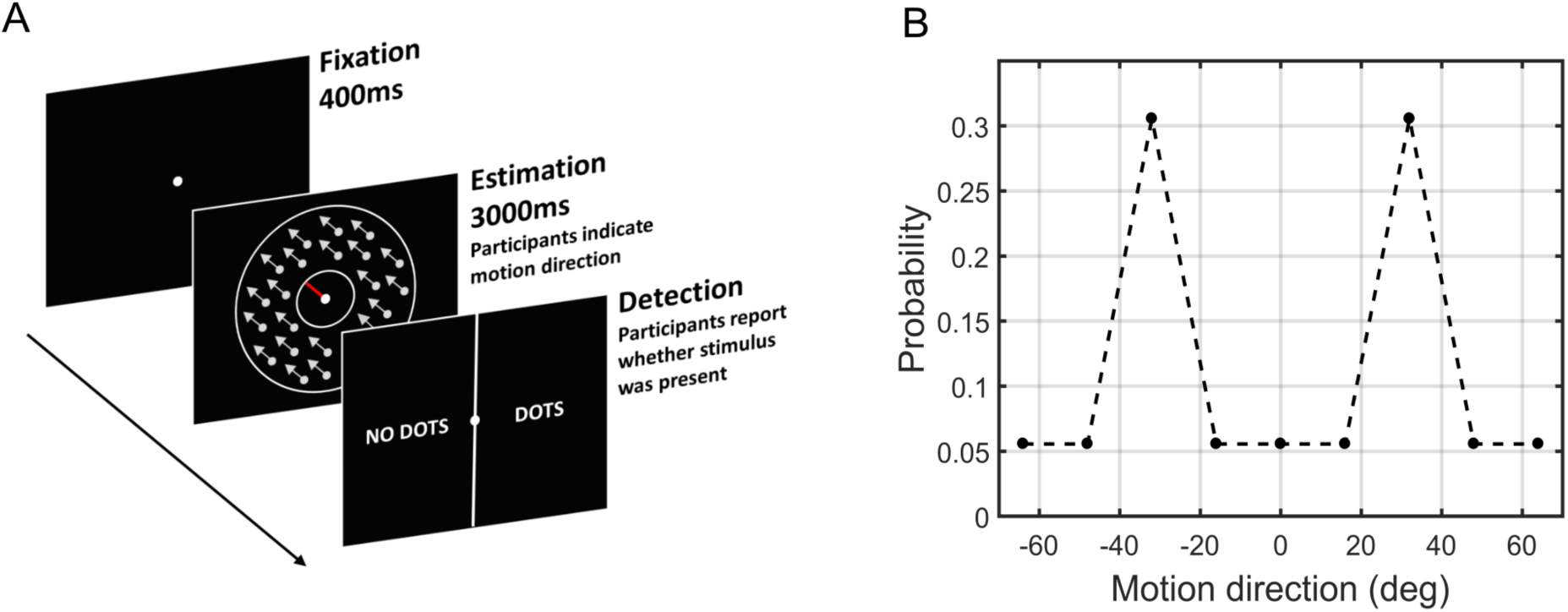
**(A)** Experimental procedure. Participants were presented with a fixation point followed by the motion stimulus and a response bar (red bar) that they were instructed to align to the perceived motion-direction. The screen was cleared either when participants clicked to validate their estimation or 3000 ms had elapsed. A new screen appeared with a two-alternative forced choice task (2-AFC), requiring participants to indicate whether they perceived the dots during the estimation task. **(B)** Probability distribution of the motion directions. Unbeknownst to participants, the distribution of motion direction was bimodal (i.e. stimuli appeared most often at ±32° from a central direction). The central direction was randomised for each participant.

### Design

Participants completed 567 trials (i.e. lasting approximately 40 minutes *vs*. 850 trials lasting 60 minutes in Chalk *et al*. 2010) with opportunities for breaks every 170 trials to prevent fatigue. Stimuli were presented at four different randomly interleaved contrast levels. The highest contrast level was 1.7 cd/m^2^ above the 5.2 cd/m^2^ background. There were 167 trials at zero contrast and 67 trials at high contrast. Contrasts of other stimuli were determined using a 4/1 and 2/1 staircase on detection performance (García-Pérez, 1998). Throughout the experiment, there were 90 trials with the 2/1 staircase and 243 trials with the 4/1 staircase. On a given trial, the direction of motion for the two staircased contrast levels could either be 0°, ±16°, ±32°, ±48°, and ±64° with respect to a central reference angle. This central reference angle was randomised for each participant.

Unbeknownst to participants, we manipulated their expectations about which motion directions were most likely to occur by presenting stimuli moving at ±32° more frequently (resulting in a bimodal distribution, **Fig. 1B**). At the highest contrast level, 50% of trials were at ±32° and 50% remaining trials at random directions (i.e. not just the predetermined directions).

### Behavioural data analysis

Performance on high contrast trials was used as an indicator of whether participants were performing the task adequately. Detection accuracy of at least 70% and estimation root mean square error (RMSE) of less than 30 degrees were the minimum criteria. All 23 controls met these criteria, while 5 out of 25 patients did not meet at least one of the criteria and thus were excluded from further data analysis (**Supplementary Fig. 1**).

The main data analysis was performed on 2/1 and 4/1 staircased contrast levels and only on confirmed trials (i.e. trials where participants validated their choice with a click within 3000 ms in the estimation task and reported seeing dots in the subsequent detection task). The first 100 trials were excluded from the analysis to allow the staircases to converge to stable contrast levels (**Supplementary Fig. 2**). After removing these trials, the luminance levels achieved by the 2/1 and 4/1 staircases were found to be overlapping (**Supplementary Fig. 2**), thus they were combined for all further analysis. Finally, since the distribution of presented directions was symmetrical around a central reference angle, the behavioural measures at equal absolute distance from the reference angle were averaged together.

In the estimation task, the variance of participants’ direction estimates was large. As in previous work (Chalk *et al*., 2010; Gekas *et al*., 2013), we hypothesized that this variability resulted from random estimations on a proportion of trials, thus increasing substantially the variance of motion-direction estimates. To account for this, we fitted the individual estimation responses to the following distribution:

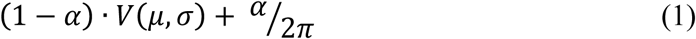

where *α* is the proportion of estimations at random directions and *V(µ*, σ) is the circular normal (i.e. von Mises; Mardia, 1972) distribution with mean *µ* and width σ.

In trials where no stimulus was presented, we reconstructed the probability distributions of participants’ responses over motion directions using Kernel Density Estimation (Silverman, 1986; Wand, 1994). The KDE is a non-parametric method used to estimate the probability density function from discrete measures of a continuous variable. To do so, a kernel that defines the form of the probability density function (e.g. von Mises kernel) was placed at each of the observed measurement. Then, all the individual kernels were summed to create the probability density function of the random variable (motion direction).

To assess the effects of the acquired expectations in both groups and to simultaneously determine whether patients differed from controls, we performed 2-way ANOVA with motion direction as a within-subjects factor and group as a between-subjects factor; and Bonferroni post hoc tests for all pairwise comparisons. All analysis was done using SPSS version 24. To determine the strength of the evidence for the null hypothesis (i.e. finding no group difference vs. finding evidence that the groups are similar) we also report Bayes factors using the Bayesian statistical software package JASP. We report Kendall correlation coefficients whenever the data is ordinal with rank ties and/or strong outliers.

### Modelling

To test for individual variability in the underlying perceptual inference and to obtain more direct measurements of the acquired expectations, we fitted a range of models to our data. The first class of models assumed that the biases were of a perceptual nature, as conceived in the Bayesian framework: sensory information is combined with a learned prior of the stimulus statistics in a probabilistic way. The simple ‘BAYES’ model assumed that the likelihood precision was constrained to be the same across all presented motion directions (corresponding to the hypothesis that there was no learning in the likelihood due to the distribution of the motion directions). An additional variant of the ‘BAYES’ model tested the hypothesis that lapse estimations were not completely random, but instead were made according to the acquired prior expectations. We call these responses ‘prior-based lapses’ (**Fig. 4**). This model was termed ‘BAYES_P’ and was otherwise equivalent to ‘BAYES’.

**Figure 2:**
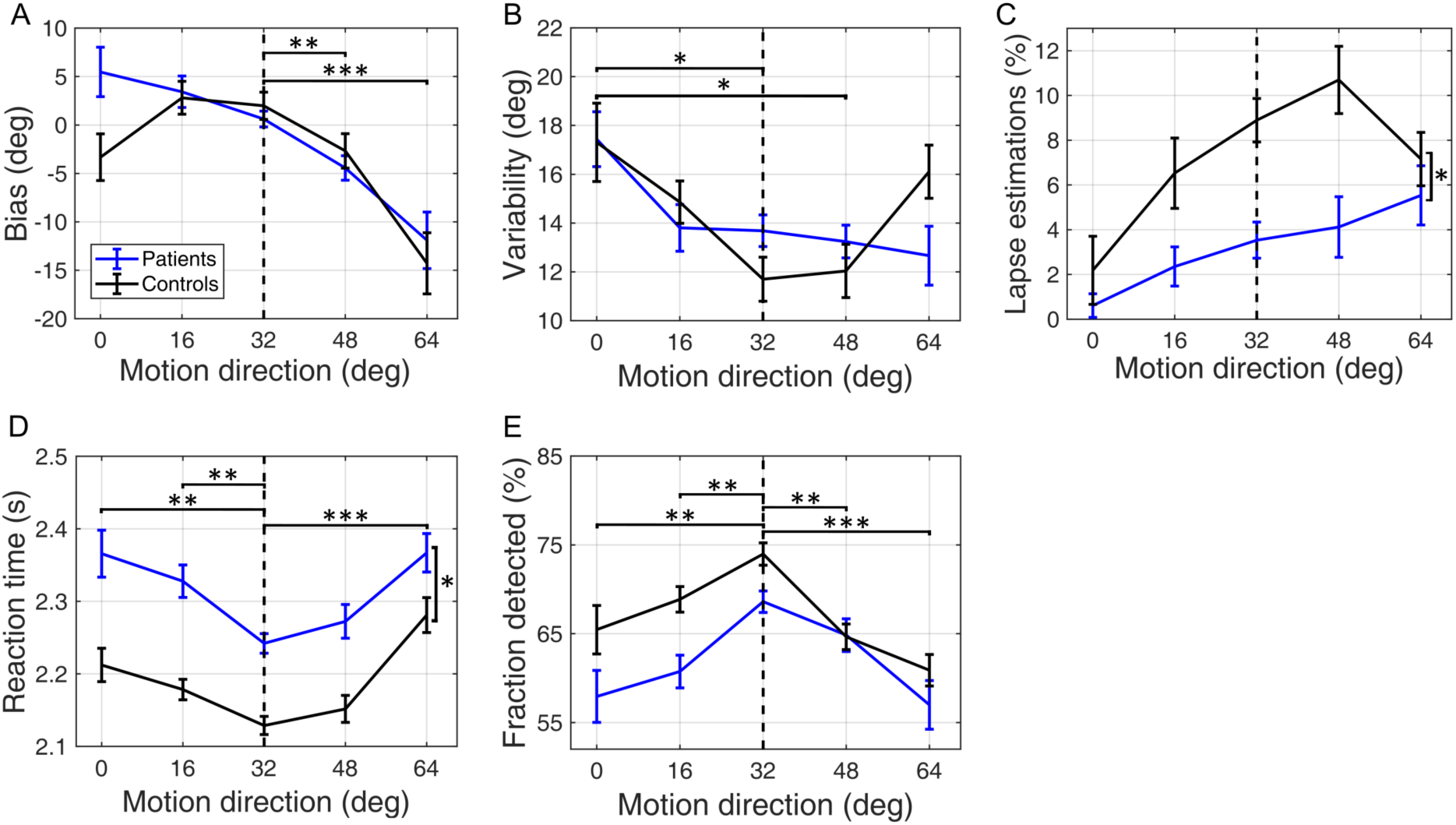
Performance on low contrast trials by patients (blue lines) and controls (black lines). **(A)** Mean estimation bias as a function of the true motion direction. **(B)** Estimation standard deviation (i.e. variability) as a function of the presented of motion direction. **(C)** Lapse estimations as a function of motion direction, estimated using (eq. 1). **(D)** Reaction times during the estimation task as a function of motion direction. **(E)** The fraction of trials in which the stimulus was detected as a function of the presented motion direction. The error bars represent within-subject standard error. The vertical dashed lines correspond to the most frequently presented motion directions (i.e. ±32°). *, ** and *** indicate significance levels at p<0.05, p<0.01, p<0.001 respectively.

**Figure 3.**
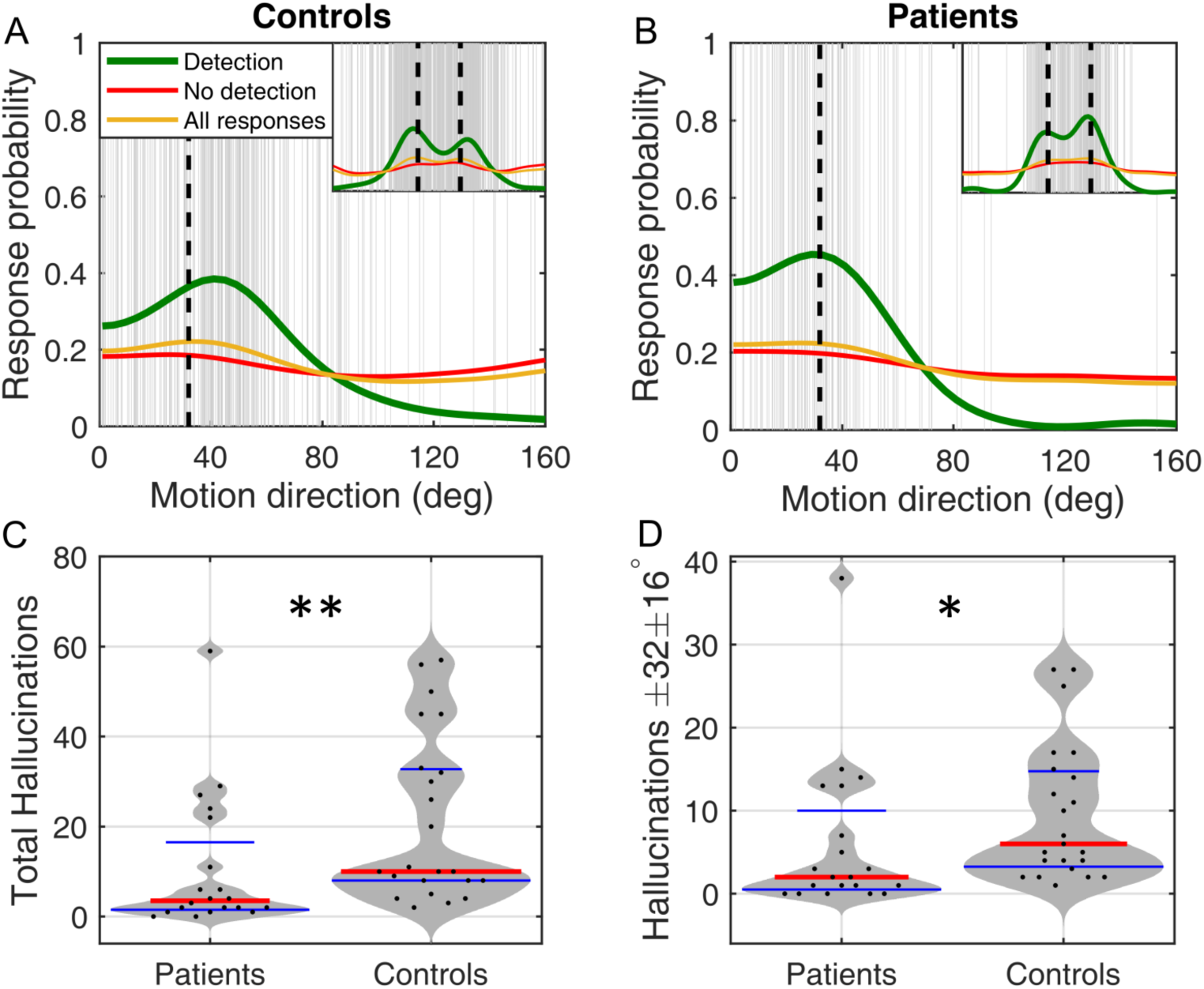
Estimation responses in the absence of stimulus. **(A, B)** Distribution of the estimation responses by patients and controls, respectively. The vertical grey lines represent reported motion directions when no stimulus was present (i.e. hallucinations) pooled across each group. The green line denotes the probability distribution of these hallucinations, which was produced using Kernel Density Estimation. The red line denotes probability distribution of responses that were followed by participants’ reporting seeing no stimulus. The orange line denotes all estimations regardless of the detection response. In the main plots the data is averaged across the central motion direction, while the insets show the corresponding distributions across the full range. **(C, D)** Comparison of patients and controls by **(C)** the total number of hallucinations (p = 0.004, two-sided rank-sum test) and **(D)** the number of hallucinations around the most frequently presented motion directions (within ±16° of ±32°; p = 0.016, two-sided rank-sum test). Red horizontal lines denote median values; blue horizontal lines denote 25th and 75th percentiles. Black dots denote individual participants, grey areas represent density of the data points. * and ** indicate significance levels at p< 0.05 and p<0.01 respectively.

**Figure 4.**
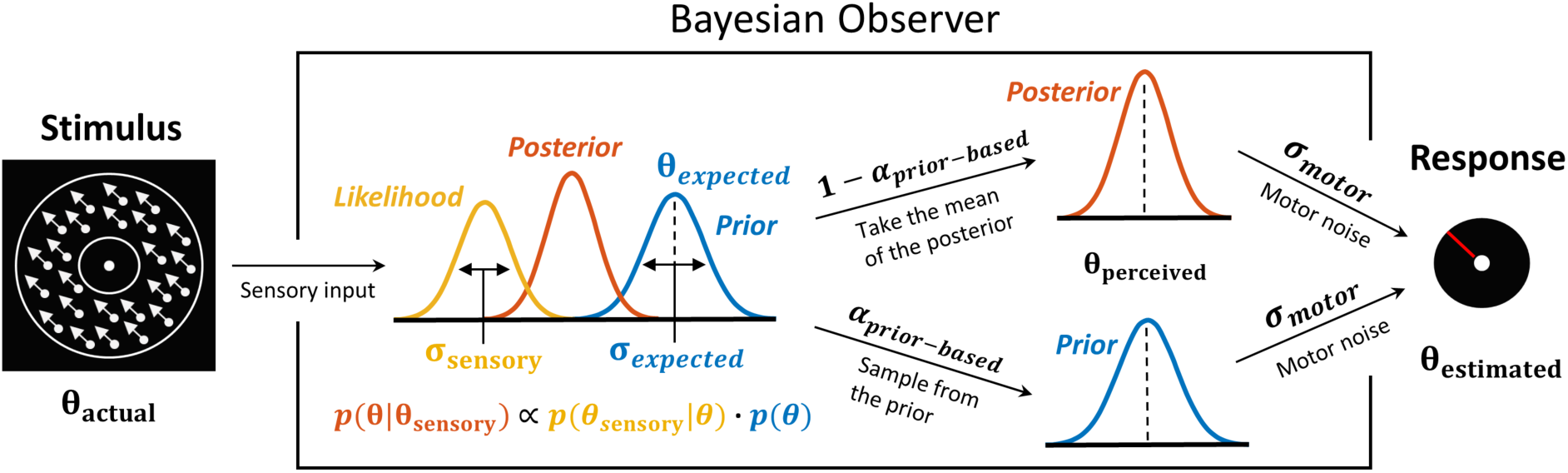
Bayesian model of estimation response for a single trial. The actual motion direction (θ_actual_) is corrupted by sensory uncertainty (σ_sensory_), and then combined with prior expectations (mean θ_expected_ and uncertainty σ_expected_) to form a posterior distribution. The perceived motion direction (θ_perceived_) then corresponds to the mean of the posterior distribution. However, on a fraction of trials, determined by the lapse rate (α_prior-based_), the perceived motion direction is sampled form the prior. Finally, in both cases, the response (θ_estimated_) is made by perturbing θ_perceived_ with motor noise (σ_motor_). This results in 4 free model parameters: σ_sensory_, σ_expected_, θ_expected_ and α_prior-based_. The motor noise (σ_motor_) is estimated from high contrast trials and is used as a fixed parameter.

Another class of models assumed that task performance could be explained by response strategies that do not involve Bayesian integration (heuristic models). According to these models, on any given trial participants responses were based solely on either the prior expectations or sensory information. We considered four variations of response strategy models (see Methods and Supplementary information for details). Below we present the Bayesian models as they provided a better explanation to the data (see Figure 5, model comparison).

**Figure 5.**
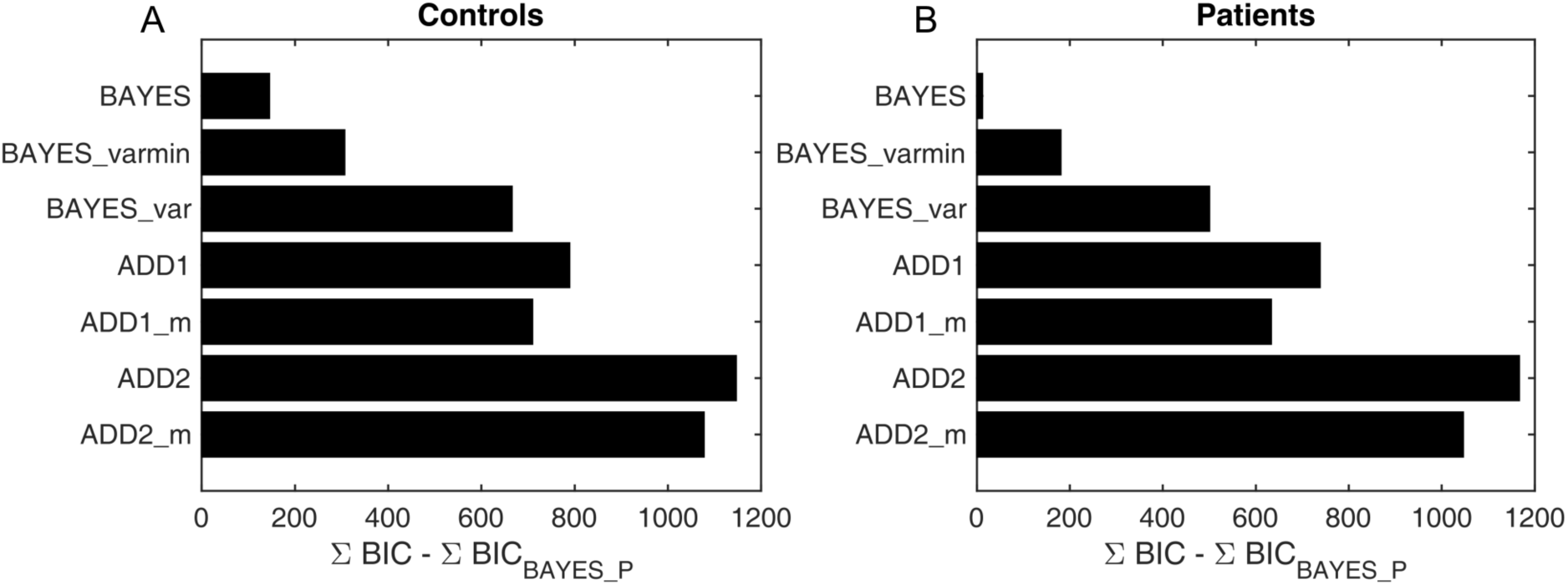
Model comparison for using Bayesian Information Criterion (BIC). **(A)** controls, **(B)** patients. X-axis measures the relative difference between BIC of each model (as indicated on Y-axis) and BIC of BAYES_P (winning model) summed across participants. Smaller BIC indicate a better model fit while being penalized for added model complexity. For both patients and controls BAYES_P provided the best model evidence, 12 BIC units better than BAYES for patients and 146 BIC than BAYES for controls. A difference of 10 units or more is considered overwhelming or decisive evidence in favor of the winning model when compared to others (Kaas & Raftery, 1995).

### Bayesian model

Following the Bayesian framework, we assumed that participants combined sensory information (likelihood) with their expectations about the motion direction (prior) on every trial. The sensory likelihood of the observed motion direction (θ_sensory_) was parameterised as a von Mises circular normal distribution with variance σ_sensory_:

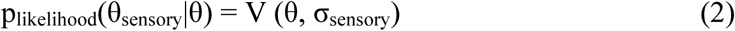

The mean of this distribution depended on the actual presented motion direction (θ_actual_), and to account for trial-to-trial variability it was drawn from another von Mises distribution centred on θ_actual_ with variance σ_sensory_ (i.e. V (θ_actual_, σ_sensory_)).

We then hypothesized that participants acquire an approximation of the ‘true’ prior from experience (p_prior_), representing the participants’ expectations of motion directions. The acquired priors were parameterized as the sum of two von Mises circular normal distributions, centred on motion directions θ_expected_ and -θ_expected_, each with variance σ_expected_:

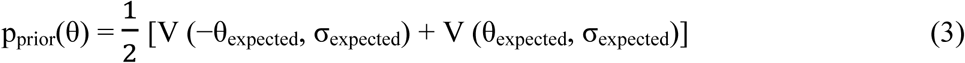

Combining the prior and the likelihood gives us the posterior probability that the stimulus is moving in a direction θ:

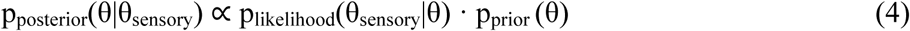

The perceived direction, θ_perceived_, was taken to be the mean of the posterior distribution (almost identical results would be obtained by using the maximum instead).

Finally, we accounted for motor noise (i.e. aligning and clicking the mouse) and lapse estimations, such that:

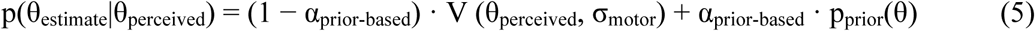

where σ_motor_ is the motor noise and α_prior-based_ is the proportion of prior-based lapse estimations on each trial (i.e. lapse estimations that follow the participants’ acquired expectations – p_prior_(θ)). That is we posited that when participants are shown a stimulus below their contrast threshold, it effectively becomes a no-stimulus trial, where participants might ‘hallucinate’ a stimulus at the most expected motion directions, sampling from their prior distribution (Laquitaine and Gardner 2018, see Fig. 7). We called this model ‘Bayes_P’ for Bayes with Prior-based lapses.

**Figure 6.**
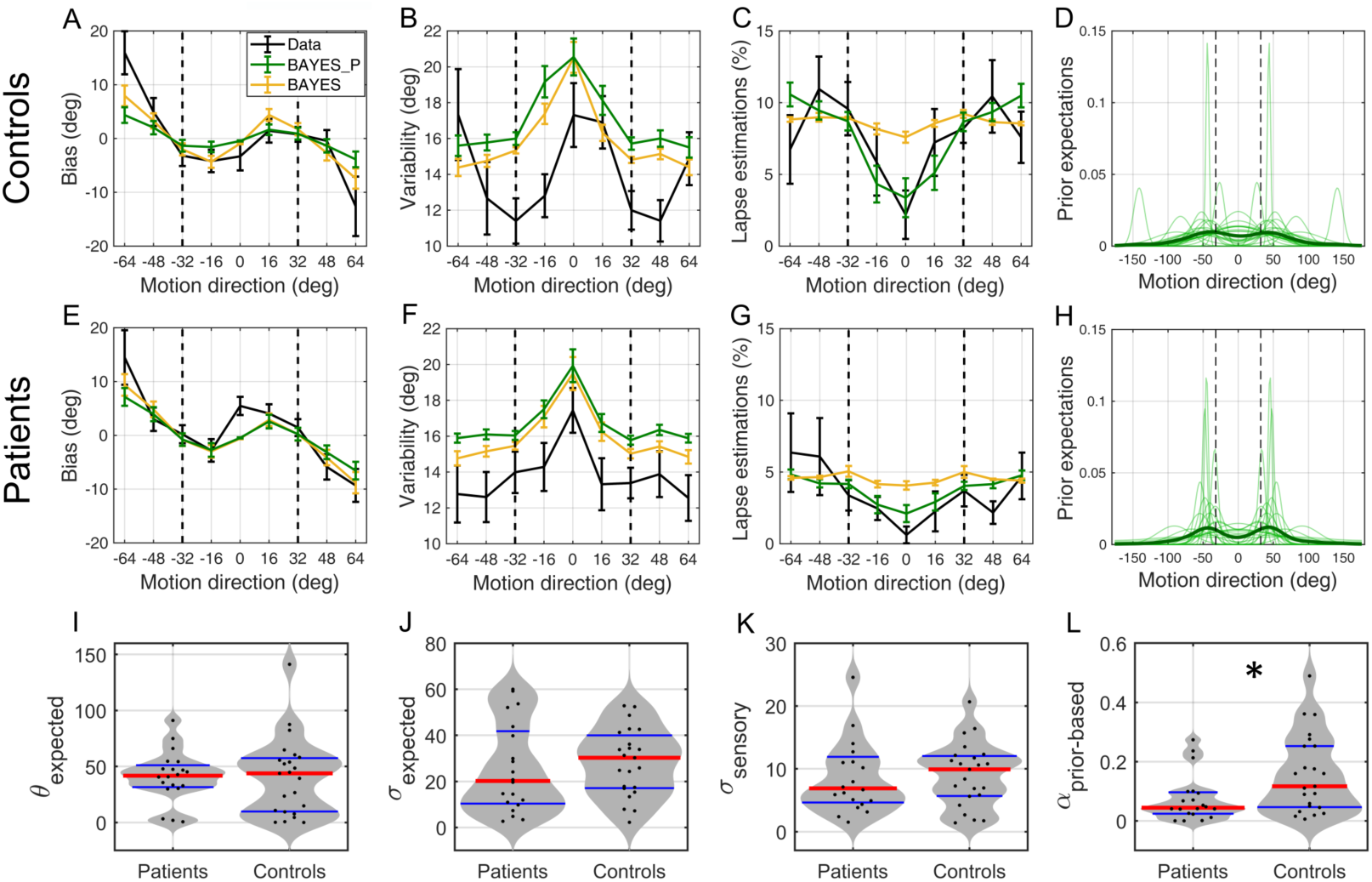
Model fits and parameter estimates. **(A-H)** Model fits for the best fitting model BAYES_P (green) and the second best model BAYES (yellow), to the behavioral data (black). **(A-D)** controls and **(E-H)** patients. **(A, E)** estimation bias, **(B, F)** estimation variability, **(C, G)** estimation lapse rate, **(D, H)** prior expectations of each individual (transparent green) and group average (thick green) as estimated via BAYES_P model. The vertical dashed lines correspond to the most frequently presented motion directions (i.e. ±32°). The error bars represent within-subject standard error. **(I-L)** Comparison of BAYES_P model parameter estimates of patients and controls. **(I)** θ_expected_ – the mean of acquired prior (p = 0.874, two-tailed rank-sum test; BF01 = 3.32), **(J)** σ_expected_ – the uncertainty in the acquired prior (p = 0.401, two-tailed rank-sum test; BF01 = 2.95), **(K)** σ_sensory_ – the uncertainty of sensory likelihood (p = 0.742, two-tailed rank-sum test; BF01 = 2.96), **(L)** α_prior-based_ – prior-based lapse rate (p = 0.024, two-tailed rank-sum test). Red horizontal lines denote median values; blue horizontal lines denote 25th and 75th percentiles. Black dots denote individual participants, grey areas represent density of the data points. * indicates significance level at p< 0.05.

**Figure 7.**
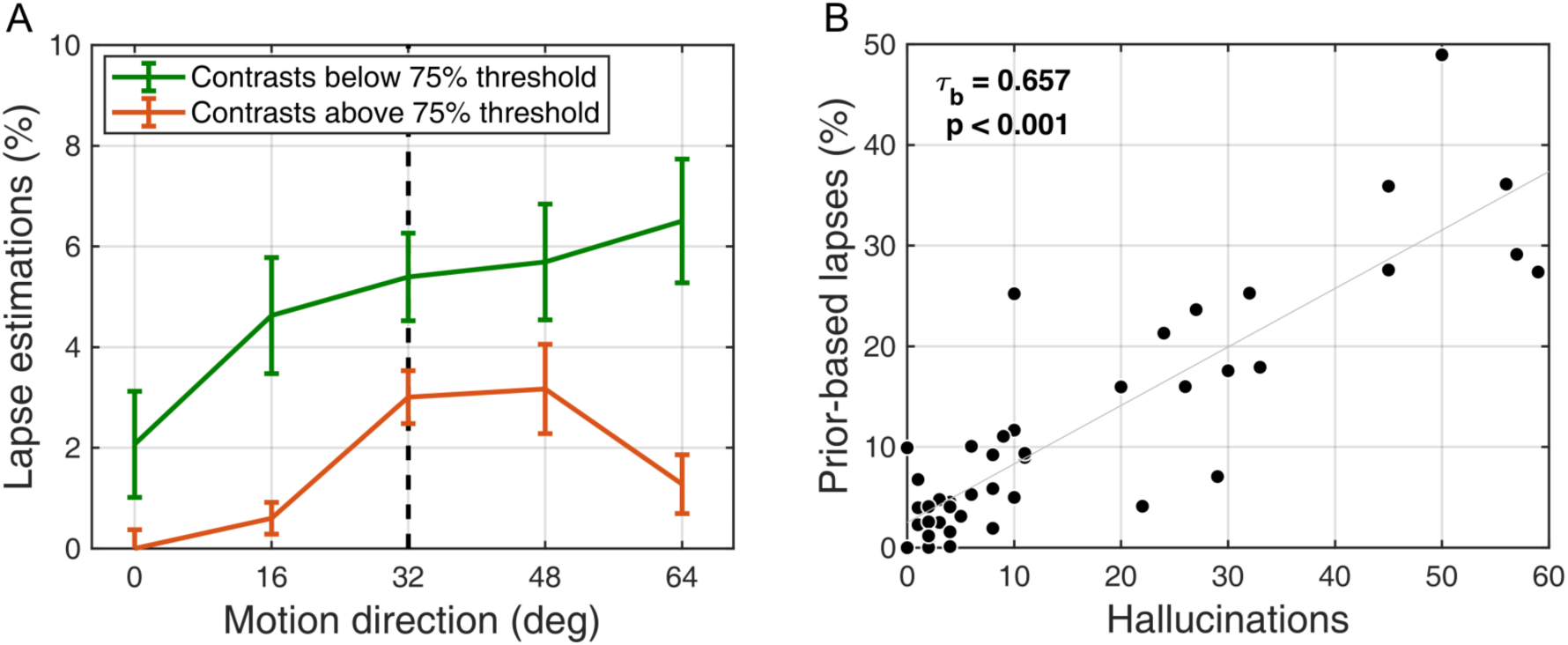
Relationship between lapse estimations and hallucinations. (**A**) The amount of lapse estimations at different stimulus contrast levels. Contrast staircase trials were split into two subsets along the 75% detection threshold, which was determined for each individual from their psychometric curves. The data then was pooled across both patient and control groups. We found that the amount of lapse estimations depended on the presented contrast level with more lapse estimations being made at lower contrasts (F(1,84) = 12.61, p < 0.001), which supported our interpretation that these estimations were not merely attentional lapses but hallucinations on trials when stimulus was hard to detect. The error bars represent within-subject errors. (**B**) Prior-based lapses and hallucinations. A strong positive correlation between prior-based lapse estimations (recovered via BAYES_P model) on low contrast trials and hallucinations on no-stimulus trials (τ_b_ = 0.657, p < 0.001; Kendall’s correlation) provided further support that both of these behaviours are driven by the same mechanism (i.e. both being samples from the prior).

Finally, we also tested variants of this model, where lapse estimations were uniformly distributed, rather than following the participants expectations (model ‘BAYES’), or that due to increased exposure to stimuli at specific angles, sensory uncertainty σ_sensory_ could vary across angles (0°, ±16°, ±32°, ±48°, ±64°; model ‘BAYES_var’), or that sensory uncertainty varied only at the most presented directions (model ‘BAYES_varmin’ – see Methods and Supplementary information for details).

## Results

### Detection performances and contrast levels

Participants’ detection performance was monitored to adapt the stimulus contrast to each participant’s just noticeable difference (JND, a measure of contrast sensitivity). Using 2/1 and 4/1 staircases, we ensured that the individual detection performances would converge to 70.4% and 84.1% respectively (Levitt, 1971).

Contrast staircases converged to stable luminance levels after about 100 trials for both groups (**Supplementary Fig. 2**); Controls converged to 0.41 cd/m^2^ (±0.03) for 2/1 staircase and 0.46 cd/m^2^ (±0.03) for 2/1 staircase while patients converged to 0.57 cd/m^2^ (±0.04) for 2/1 staircase and 0.62 cd/m^2^ (±0.05) for 4/1 staircase. These results confirm previous findings (Skottun and Skoyles, 2007) suggesting that patients with schizophrenia display significantly poorer contrast-sensitivity in comparison to controls (2/1 staircase: Z = 3.15, p = 0.002; 4/1 staircase: Z =2.90, p = 0.004, two-tailed rank-sum test).

### Statistical learning

First, we investigated whether participants acquired the statistics of the stimulus. To do so, we looked at patterns suggestive of statistical learning in each group, namely attractive biases towards the most frequent directions, decreased reaction times and improved detection performance for the most frequent directions (**Fig. 2**).

### Estimation performance

To investigate whether the participants’ perceived motion-directions were biased, we measured the difference between the true motion direction and the motion direction reported by the participants. **Fig. 2A** displays the average estimation bias plotted against the true motion direction for each group. Overall, there was a significant effect of motion direction on the estimation bias (F(2.45, 100.52) = 15.37, p < 0.001, 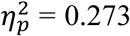, Greenhouse-Geisser correction ε = 0.613), but no differences between the groups (group main effect: F(1, 41) = 0.83, p = 0.369, 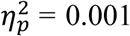; with moderate to substantial evidence for the null hypothesis, BF_01_ = 3.99); and no group*angle interaction (F(2.45, 100.52) = 1.64, p = 0.193, 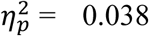). Pairwise comparisons (with Bonferroni correction) revealed that there was an attractive bias towards ±32° at ±48° and ±64° (MD = 4.858, p = 0.002; and MD = 14.395, p < 0.001, respectively), but not at ±16° (MD = 1.818, p = 0.955). Together, these results confirm that both patients and controls were biased towards perceiving motion directions as being more similar to the most frequently presented directions than they really were, consistent with having acquired the priors that approximate the statistics of the stimulus.

We also investigated whether the effects of acquired prior expectations were reflected in the variability of estimations, namely a decrease of variability for the expected directions (**Fig. 2B**). We found a significant main effect of motion direction (F(3.07, 125.69) = 5.18, p = 0.002, 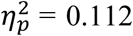, Greenhouse-Geisser correction ε = 0.766), but no differences between the groups (main effect of group: F(1, 41) = 0.02, p = 0.880, 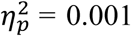; with moderate to substantial evidence for the null hypothesis, BF_01_ = 3.74); and no group*angle interaction F(3.07, 125.69) = 1.58, p = 0.196, 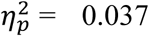). Pairwise comparisons showed that variability at 0° stood out the most, being significantly larger than at ±32° and ±48° (MD = 4.680, p = 0.012; and MD = 4.733, p = 0.025, respectively), although not different than at ±16° and ±64° (MD = 3.044, p = 0.239 and MD = 2.990, p = 0.541). The increased variability in the region between the two modes reflects their conflicting influence on the percepts in this region.

Finally, we analysed lapse estimations, which were captured by the ‘α’ term in Eq. (1) and which were assumed to arise from random responses on some of the trials (**Fig. 2C**). We found both motion direction and group main effects to be significant (F(4,164) = 5.76, p < 0.001, 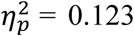 and F(1, 41) = 6.41, p = 0.015, 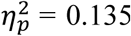, respectively), with patients exhibiting fewer lapse estimations. Pairwise comparisons revealed that lapse rate at 0° was significantly smaller than at all other directions (±32°, MD = 4.814, p = 0.001; ±48°, MD = 6.010, p = 0.007; ±64°, MD = 4.951, p = 0.003), except for ±16° (MD = 3.043, p = 0.393). The finding that the estimated lapses would depend on the presented motion direction was surprising: if the lapses were completely random, we should observe no such dependence in the estimated values. We therefore postulated that when participants make guesses about the direction of motions, their estimations might be sampled from the distribution of the acquired prior expectations rather than uniformly. Later, we explicitly tested this possibility via computational modelling and found that it indeed was the case (see Modelling results).

### Reaction times and Detection Performance

Next, we examined how participants’ acquired expectations influenced reaction times and the detection of stimulus. The estimation reaction times **(Fig. 2D**) show a significant main effect of motion direction (F(2.73,111.76) = 10.80, p < 0.001 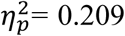, Greenhouse-Geisser correction ε = 0.681). This was driven by decreased reaction times at the most frequent directions as revealed by pairwise comparisons: reaction time at ±32° was significantly shorter than at all other directions (0°, MD = 0.104, p = 0.001; ±16°, MD = 0.068, p = 0.004; ±64°, MD = 0.139, p < 0.001), except for ±48° (MD = 0.027, p = 1.000). Furthermore, patients were found to also be significantly slower than controls (F(1, 41) = 4.11, p = 0.049, 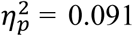), but we found no interaction between group and motion direction (F(2.73, 111.76) = 0.66, p = 0.563, 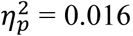).

Slow reaction time is a hallmark of schizophrenia that has been documented thoroughly in the literature in simple reaction-time tasks using visual and/or auditory stimuli (e.g., see Nuechterlein, 1977; Fioravanti *et al*., 2012).

An even more direct way of assessing how the acquired expectations influenced the detection of stimulus is to analyse the fraction of trials where participants explicitly report seeing or not seeing the stimulus (**Fig. 2E**). We found that the detection of stimulus was greatly affected by the presented motion direction (F(2.36,96.64) = 8.51, p < 0.001, 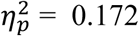, Greenhouse-Geisser correction ε = 0.589), with stimulus at ±32° being the most frequently detected direction as shown by pairwise comparisons: detection at ±32° was significantly better than at all other directions (0°, MD = 9.59, p = 0.004; ±16°, MD = 6.48, p = 0.001; ±48°, MD = 6.54, p = 0.001; ±64°, MD = 12.35, p < 0.001). However, the groups were not found to be different (main group effect: F(1,41) = 3.62, p = 0.064, 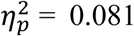; although there was no evidence for the null hypothesis either, BF_01_ = 0.97); group*motion direction interaction was also non-significant: F(2.36,96.64) = 1.11, p = 0.340, 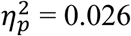).

Overall, these results indicate that, in terms of detection responses (hit rates and reaction-time), similar benefits of statistical learning were present in both patient and control groups. Overall, behavioural measures suggest that prior effects (e.g. Bias, RT, and Hit rate) became significant as early as within first ∼100 trials for both patients and controls (see Supplementary Figures 3-4).

### Perceived motion in absence of visual stimuli (hallucinations)

Finally, we investigated whether the acquired statistics about the motion stimulus affected the participants’ perception on trials where no stimulus was presented, but where participants reported both a motion direction and seeing a stimulus. We refer to this effect as hallucinations. These hallucinations in our perceptual task are of course different in terms of content and complexity from the visual hallucinations observed in psychosis. However, the underlying mechanisms might be informative to the understanding of illusions and hallucinations in schizophrenia (Silverstein and Keane 2011a; 2011b; Notredame *et al*., 2014).

To quantify the probability ratio that participants made estimates that were closer to the most frequently presented motion directions relative to other directions, we multiplied the probability that participants estimated within 16° of these motion-directions by the total number of 32° bins:

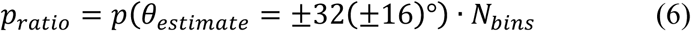

This probability would be equal to 1 if participants were equally likely to estimate within 16° of ±32° as they are to estimate within the other 16° bins.

We found that the median value of *‘p*_*ratio*_*’* was significantly greater than 1 for both patients and controls (median(*p*_*ratio*_) = 2.88, p = 0.003 and median(*p*_*ratio*_) = 2.75, p <0.001, respectively; two-tailed signed-rank test), indicating that both patients and controls were much more likely to hallucinate the most frequent motion directions as opposed to all other directions (**Fig. 3A, B**). Bayesian statistical analysis provided moderate to substantial evidence for the groups being the same in this measure (BF_01_ = 3.32). Finally, hallucinations of the most frequent directions (i.e. hallucinations at ±32°±16°) were quick to develop during the course of the experiment: they became significant after only 150 trials for both controls (*p=0*.*036*, one-tailed signed-rank test; Supplementary Fig. 3D) and patients (*p=0*.*035*, one-tailed signed-rank test; Supplementary Fig. 4D).

While both patients and controls hallucinated predominantly towards the most frequently presented directions (i.e. prior-based hallucinations), patients exhibited fewer of such hallucinations (**Fig. 3C**; hallucinations within ±16° of ±32°; p = 0.016, two-sided rank-sum test), and also exhibited less hallucinations overall (**Fig. 3D**; p = 0.004, two-sided rank-sum test). We wanted to know whether the severity of the symptoms was predictive of the magnitude of this effect. However, we found no correlations between the number of hallucinations and the PANSS positive, negative, general or total scores nor duration of illness, or between the daily-dosage of anti-psychotics (Olanzapine equivalent; Leucht et al., 2015) and the total number of hallucinations.

We also investigated the distribution of responses when participants estimated the direction of motion but reported not seeing any dots. Interestingly, in this subset of trials, and unlike in our previous work (Chalk et al, 2010), estimations were also more likely than chance to be made around the most frequent motion directions by both patients and controls (median(*p*_*ratio*_) = 1.24, p = 0.002 and median(*p*_*ratio*_) = 1.41, p = 0.045, respectively; two-sided signed rank test). One explanation for this might be habitual effects - when the bar is moved towards the most frequent directions out of motor habit. Another possible explanation is that participants might have hallucinated stimuli in the expected prior directions, but their confidence about their percept being very low, they sometimes chose to report not seeing any dots in the hope to give the correct answer (each detection response was followed by immediate feedback).

### Modelling results

We fitted the models to the behavioral data and computed the Bayesian Information Criterion (BIC) for each participant, which gives us information regarding model fit while penalizing for extra model complexity (preventing overfitting). We found that the BAYES_P model had the smallest BIC for both patients and controls (**Fig. 5**) – indicating best performance, with a difference in BIC between the winning model (BAYES_P) and the second best model (BAYES) being larger than 10. This is equivalent to a log Bayes factor larger than 10, and is considered to be very strong or ‘decisive’ evidence in favor of the winning model (Kaas & Raftery, 1995). Model fits to the data also showed that BAYES_P was much better at fitting lapse estimations, confirming that such estimations followed the acquired prior distribution instead of being random (**Fig. 6C and G**). Moreover, we found a strong correlation between the prior-based lapses recovered via BAYES_P and hallucinations (τ_b_ = 0.657, p < 0.001; Kendall’s correlation; **Fig. 7B**), suggesting that prior-based lapses could be considered as hallucinations experienced on trials when stimulus is too weak to be detected (**Fig. 7A**).

Finally, we compared patients and controls on the basis of BAYES_P parameter estimates (**Fig. 6I-L**). Consistent with the behavioral data analysis, we found no differences in the acquired prior expectations (**Fig. 6I, J;** the mean of acquired prior: p = 0.874, BF_01_ = 3.32; and the uncertainty in the acquired prior: p = 0.401; two-tailed rank-sum test; BF_01_ = 2.95). There were no differences in the precision of sensory likelihood (**Fig. 6K**, p = 0.742, two-tailed rank-sum test; BF_01_ = 2.96). Lastly, just as in the behavioral data, we found that patients made less prior-based lapse estimations (**Fig. 6L**, α_prior-based_: p = 0.024, two-tailed rank-sum test), suggesting that the model parameter ‘α_prior-based_’ captures the participants’ propensity to experience prior-based hallucinations in our task.

### Parameter recovery for model BAYES_P

To gauge the reliability of our modelling results, we performed parameter recovery for the winning BAYES_P model. Parameter recovery consists of simulating synthetic data with different sets of known parameter values (‘true parameters’) for a given model and then fitting the same model to the synthetic data to estimate and recover these parameters (‘recovered parameters’). The strength of correlation between the actual and recovered parameters measures the reliability of modelling results. Parameter recovery, just as the parameter estimation from behavioral data, is sensitive to any correlations that might be present among the model parameters, to the choice of parameter estimation methods and also to the amount of data used for model fitting. Therefore, parameter recovery serves as a crucial step in validating the reliability of the modelling results (Palminteri, et al., 2017).

We found that the winning BAYES_P model recovered parameters very well, which was reflected in the coefficient of determination (R^2^) for all recovered parameters being R^2^ ≥ 0.84 (**Fig. 8**).

**Figure 8.**
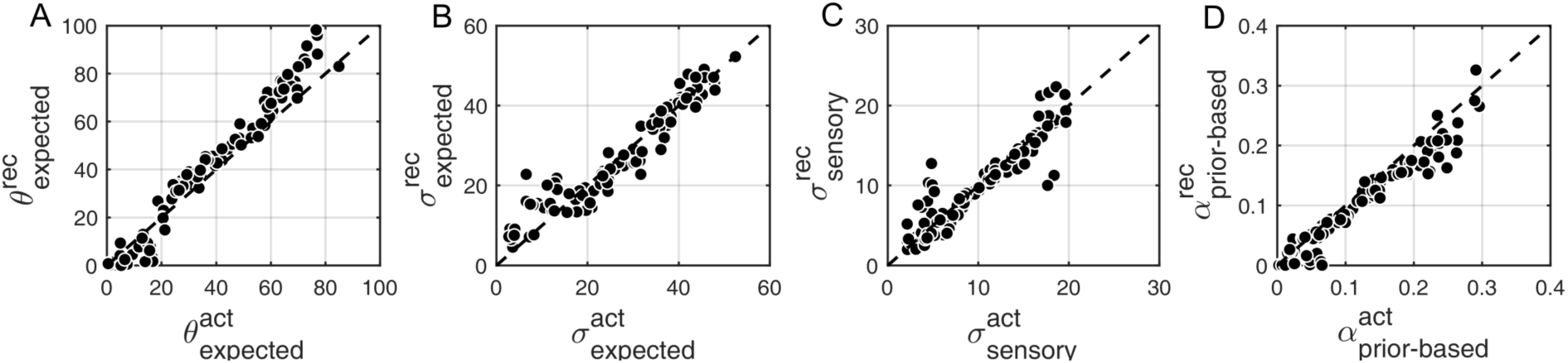
Parameter recovery with BAYES_P model. **(A)** θ_expected_ - mean of the prior expectations (R^2^ = 0.96), **(B)** σ_expected_ - uncertainty of the prior distribution (R^2^ = 0.89), **(C)** σ_sensory_ - uncertainty in the sensory likelihood (R^2^ = 0.84), **(D)** α_prior-based_ – prior-based lapse rate (R^2^ = 0.94). X-axes – actual parameters used for simulating the data (denoted with the superscript ‘act’), Y-axes – recovered parameters (denoted with the superscript ‘rec’) from fitting the model to the simulated data. The dashed diagonal line is a reference line indicating perfect parameter recovery.

## Discussion

We were interested in testing the emerging model of schizophrenia proposing that the disorder could stem from deficits in Bayesian inference (Corlett *et al*., 2009a, 2009b; Fletcher and Frith, 2009; Adams *et al*., 2013; Schmack et al., 2013, 2015, 2017; Teufel et al., 2015; Powers et al., 2017, Jardri et al., 2017). The experimental paradigm we chose is well suited to quantitatively assess the acquisition of sensory priors, how these priors are used in perception, as well as to quantify inter-individual variability in the learning and inference process (Chalk et al 2010, Karvelis et al, 2018).

### Acquisition of visual prior expectations

We found that both the control and patient groups implicitly learned the statistics of the motion stimuli and that those expectations modified their perception, consistent with them acquiring a Bayesian prior of the stimulus statistics and combining it with sensory evidence, replicating our previous results (Chalk et al, 2010). This was reflected by attractive estimation biases towards the frequently presented directions, faster reaction times and higher detection rates at these directions, as well as hallucinated motion directions in the absence of stimulus predominantly following the most frequent directions.

Patients with schizophrenia were not qualitatively, nor quantitatively different from controls in the measures used to assess learning of the task statistics. This suggest that while there are clearly domain-specific perceptual (e.g. hallucinations) and learning deficits in schizophrenia (e.g. jumping to conclusions), our study demonstrates that these deficits are not due to a domain-general impairment in the acquisition and/or utilization of statistical information in the environment. That is, we find that patients with chronic schizophrenia do not appear to be impaired in the acquisition of visual statistical priors in our task.

These results are consistent with studies finding no deficit in implicit learning in schizophrenia (Kéri *et al*., 2000; Danion *et al*., 2001; Marvel *et al*., 2005; for review see: Gold *et al*., 2009). In contrast with studies that assay explicit statistical learning and inference using more cognitive tasks (i.e. usually believed to involve frontal cortical regions), here we measured implicit statistical learning of visual stimuli that could be embodied in visual processing areas rather than frontal cortices (Kok *et al*., 2013). In fact, patients with schizophrenia appear relatively spared in implicit learning tasks that do not require integrating information after each trial (Gold *et al*., 2009). These results are also consistent with our previous study using the same paradigm showing intact statistical learning in participants with high schizotypal traits (Karvelis et al, 2018).

### Impact of acquired visual prior expectations

We found no difference between patients and controls as to the influence of the acquired expectations on their performance regarding estimation of the motion directions. They were not more or less biased towards the most frequent directions, nor more or less variable in those estimations.

Patients were found to differ from controls in three ways, however. First, patients with schizophrenia displayed significantly poorer contrast discrimination thresholds and slower reactions times, as documented in previous studies. Second, and more interestingly, patients reported significantly fewer hallucinations at all directions and fewer prior-based hallucinations (i.e. hallucinations of the most frequently presented motion directions). Third, patients exhibited fewer prior-based lapse estimations than controls on low contrast trials. Prior-based lapses can be interpreted in the same way as hallucinations on no-stimulus trials. When using contrast staircases, contrast levels hover around the detection threshold, which means that on a significant number of trials the stimulus contrast falls below the participants’ threshold of perception, effectively becoming equivalent to trials with no stimulus. If participants have hallucinations on these trials (and thus report that they have perceived a stimulus), these will be expressed in our results as prior-based lapses. In support for this interpretation, we found that there were significantly more lapse estimations made on trials when the stimulus contrast was below the 75% detection threshold then when it was above (F(1,84) = 12.61, p < 0.001; **Fig. 7A**) and we found a strong correlation between prior-based lapses and hallucinations on no-stimulus trials (**Fig. 7B**). Together, this suggests that although patients appear to acquire the same prior as controls, they tend to hallucinate this prior less than controls when there is no stimulus, or when the stimulus is below detection threshold.

The fact that patients exhibit fewer prior-based hallucinations suggests that their perception is less influenced by their learned prior expectations than observed in control participants. This is consistent with previously reported findings suggesting that patients with chronic schizophrenia are less sensitive to expectation-driven illusions (e.g. the hollow-mask illusion) than controls (Tschacher *et al*., 2006; Dima *et al*., 2009; Crawford *et al*., 2010; Horton and Silverstein, 2011; Keane *et al*., 2013, Notredame et al., 2014). This finding is also in line with results from Schmack et al. (2013, 2015), reporting a decreased influence of induced expectations (priors) on perception.

It is intriguing however that the influence of prior expectations is weaker in the detection task, but similar to that of controls for the estimation task. The absence of differences in the estimation bias could be explained by different factors.

One factor might be that very few of our patients reported clinically significant levels of hallucinations (PANSS items >3), and that this propensity might be more pertinent than the diagnosis of schizophrenia *per se* (Powers et al, 2017). We found some evidence supporting this idea: PANSS Positive symptom score and prior-based lapses (estimated via BAYES_P) were close to being negatively correlated (τ_b_=-0.314, p= 0.063; Kendall’s correlation; **Supplementary Fig. 5A**); this relationship was also backed up by a much stronger correlation for when lapse estimations were estimated directly from the behavioural data using Eq. (1) (τ_b_ = −0.465, p = 0.006; Kendall’s correlation; **Supplementary Fig. 5B**). However, it should be noted that no correlation was found between the PANSS Positive symptom score and hallucinations exhibited on no-stimulus trials.

Similarly, the absence of stronger effects might be related to illness duration: weaker estimation biases may be characteristic of earlier stages of the illness but may not be detectable anymore in our patients as they have been generally ill for a long time, due to medication or compensatory mechanisms. We find some support for this idea with a significant correlation between duration of illness and magnitude of estimation bias (τ_b_ = 0.523, p = 0.003; Kendall’s correlation; **Supplementary Fig. 5C**).

Finally, since the differences appear only for the detection part of the task, it might be that chronic patients have simply developed an increased perceptual threshold. Following this idea, patients would require stronger evidence (i.e. sharper posterior) in order to perceive a stimulus or to make a decision about the presence of a stimulus. This is consistent with the fact that patients required higher stimulus contrasts and integrated information over longer periods of time before responding (slower reaction times during the estimation task). We therefore hypothesize that it is a possible adaptation strategy used by patients over time to minimize responses to stimuli that were not truly present (i.e. their psychotic hallucinations).

Taken together, our results suggest that statistical learning is intact in relatively well patients with chronic schizophrenia on stable doses of second generation antipsychotic medication. The impact of their acquired priors is also the same as that of controls in the estimation task, but weaker in the detection task. These results are surprising in view of the current prominent theories proposing that schizophrenia is a disorder of predictive processing or Bayesian inference, and suggest ways in which these account need to be nuanced. The similarity of controls and patients’ performance in our task may however be related to the success of treatment or some other adaptive process related to illness duration. Future work will aim at testing participants at earlier stages of the illness, and ideally before pharmacological treatment has begun.

