## Supplementary Information for "Acquisition of visual priors and induced hallucinations in chronic schizophrenia"

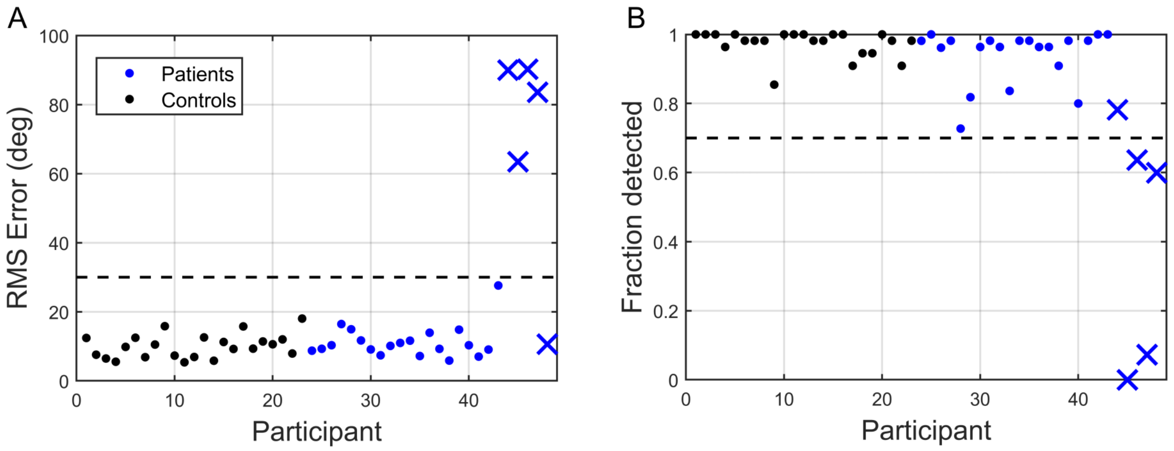


**Supplementary Figure 1:** Performance on high contrast trials. **(A)** Root mean square error of estimations. **(B)** Fraction of trials on which the stimulus was detected. The dots represent included participants; the cross marks represent excluded participants. Dashed lines denote inclusion criteria (30 degree estimation and 70% detection).

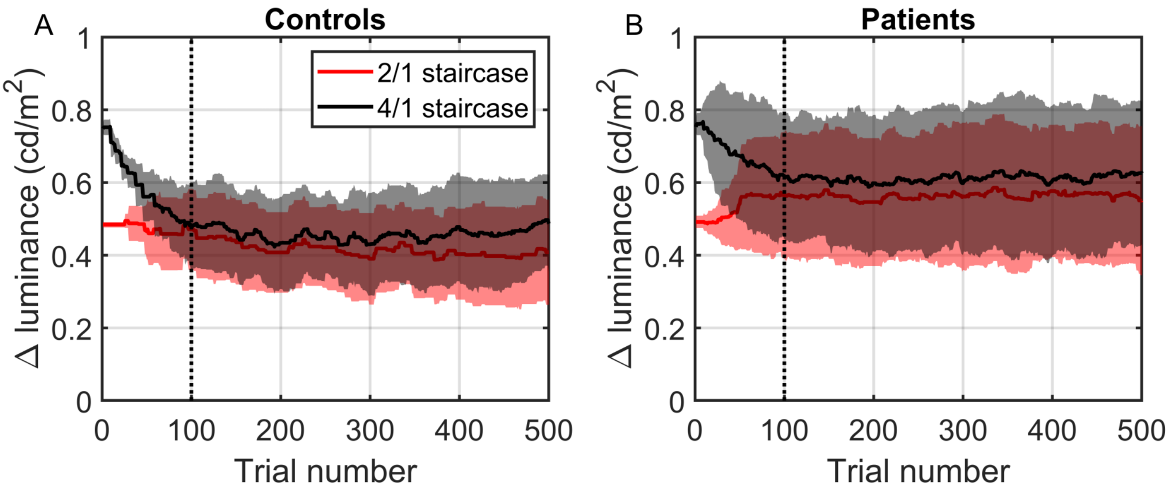


**Supplementary Figure 2:** Convergence of 2/1 and 4/1 staircase luminance levels. **(A)** Controls, **(B)** Patients. Both groups reached convergence after ~100 trials. These trials were excluded from further data analysis.

### Emergence of prior effects

We wanted to determine when the acquired expectations started to have a significant effect on performance and whether this was the same for both groups. First and most importantly, we wanted to know when the estimations on low contrast trials became biased towards ±32°. To do so, we computed cumulative moving averages at every 50 trials for the bias at ±48° with respect to bias at ±32°. Next, we looked at estimation reaction times (RT) on low contrast trials and compared mean RT of each individual at ±32° with mean RT at all other directions. Similarly, we looked at the average detection performance on low contrast trials and compared the fraction of trials in which stimulus was detected at ±32° with the mean fraction detected over all other presented directions.

We found that both patients and controls showed signs of very rapid acquisition of the priors. For the bias to become statistically significant, it took less than 100 trials for patients (**Supplementary Fig. 3A**) and less than 150 trials for controls (**Supplementary Fig. 4A**). Similarly, both patients and controls became significantly faster and significantly better at detecting stimulus moving at ±32° within the first 100 trials (**Supplementary Fig. 3B, C and 4B, C respectively**).

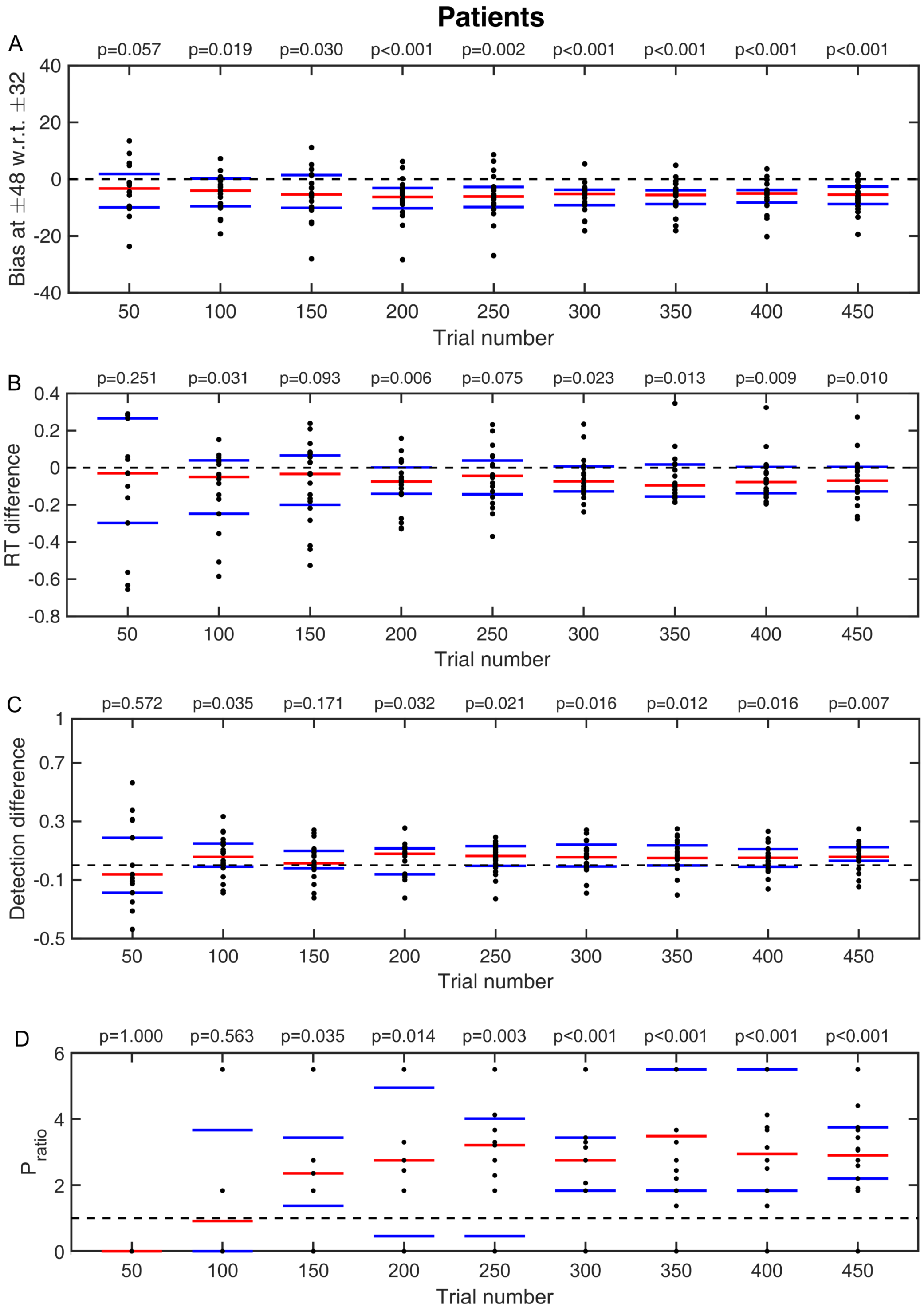


**Supplementary Figure 3:** Emergence of prior effects in patient group. **(A)** Cumulative moving averages of bias at ±48° with respect to bias at ±32°. **(B)** Cumulative moving averages of median differences between estimation RTs at ±32° and RTs at all other directions. **(C)** Cumulative moving averages of median differences between fraction of detected stimuli at ±32° and fraction detected at all other directions. **(D)** Cumulative moving averages of the probability ratio of hallucinating predominantly around ±32 on no-stimulus trials. Red bars indicate median values and blue bars indicate 25th and 75th percentiles. p-values are denoted above each plot (one-tailed Wilcoxon signed rank test).

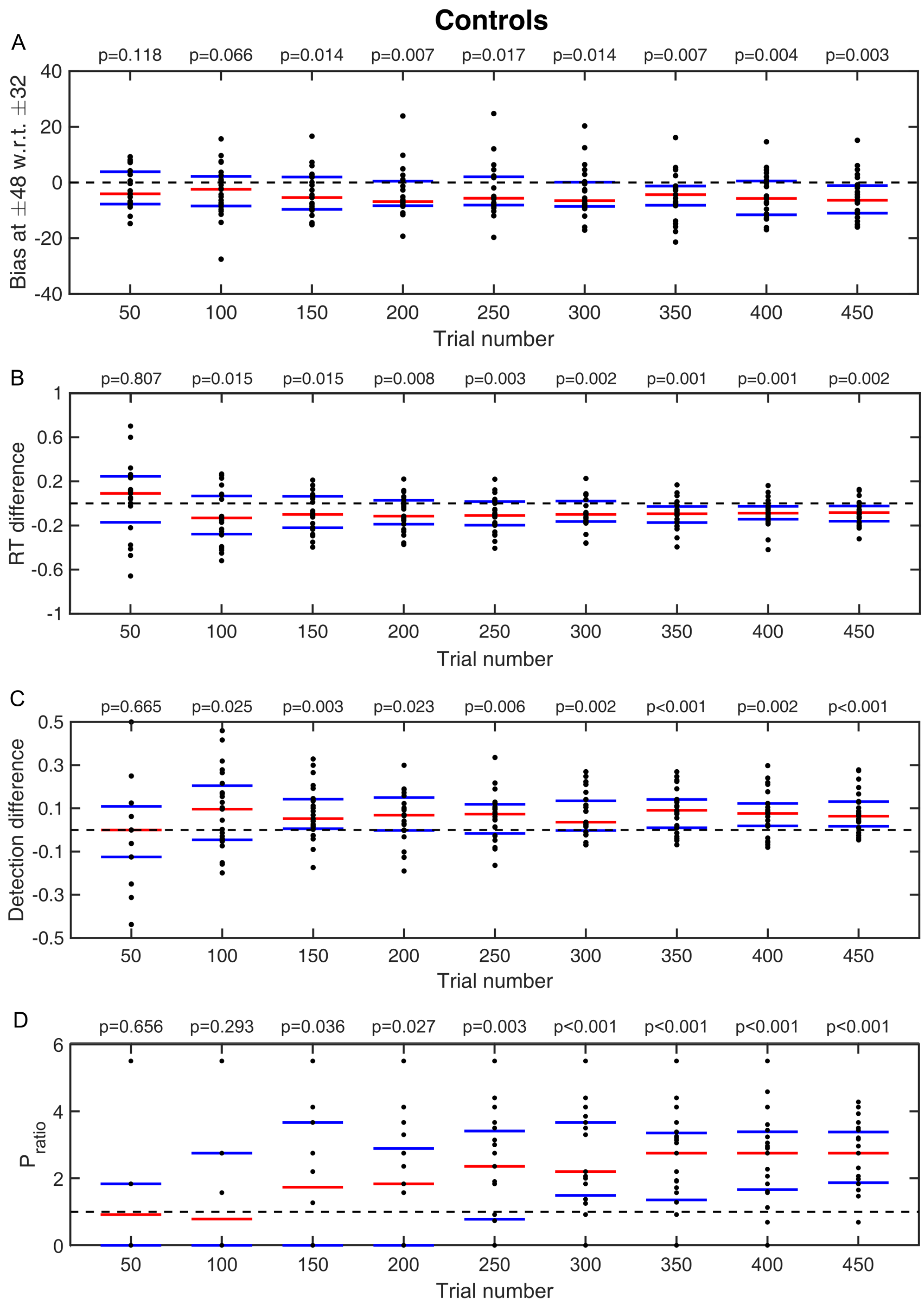


**Supplementary Figure 4:** Emergence of prior effects in control group. **(A)** Cumulative moving averages of bias at ±48° with respect to bias at ±32°. **(B)** Cumulative moving averages of median differences between estimation RTs at ±32° and RTs at all other directions. **(C)** Cumulative moving averages of median differences between fraction of detected stimuli at ±32° and fraction detected at all other directions. **(D)** Cumulative moving averages of the probability ratio of hallucinating predominantly around ±32 on no-stimulus trials. Red bars indicate median values and blue bars indicate 25th and 75th percentiles. p-values are denoted above each plot (one-tailed Wilcoxon signed rank test).

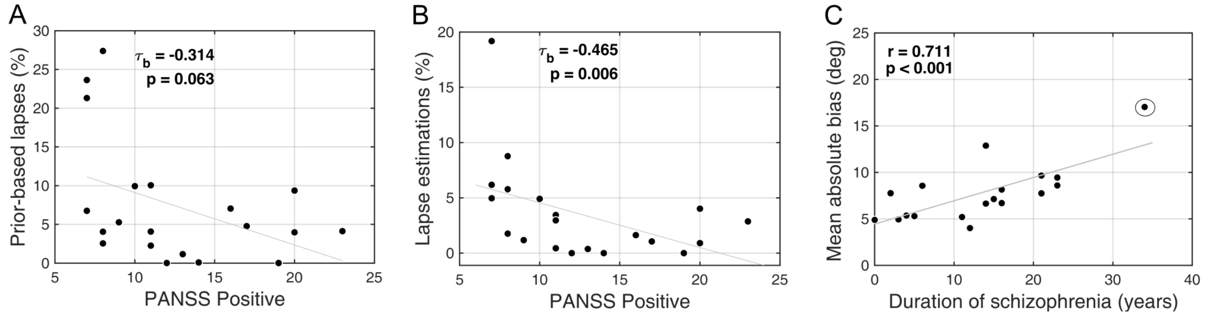


**Supplementary Figure 5:** Exploratory analysis. (**A**) Prior-based lapses (estimated via BAYES_P) as a function of PANSS positive score ($\tau_b$ = -0.314, p = 0.063; Kendall's correlation). (**B**) Lapse estimations (determined via Eq.(1)) and PANSS positive score ($\tau_b$ = -0.465, p = 0.006; Kendall's correlation). (**C**) Mean absolute bias as a function of duration of illness. The plot shows a positive relationship between the duration of illness and the amount of bias exhibited during the task (r = 0.711, p < 0.001; Pearson's correlation. The correlation remained highly significant after excluding the outlier (circled) (r = 0.523, p = 0.003; Pearson's correlation).

### Modelling

#### Variant Bayes models

The model 'Bayes' is similar to the model 'Bayes_P' described in the main text, but has uniform lapse estimations instead of 'prior-based' lapse estimations, such that:

$$p(\theta_{estimate}|\theta_{perceived}) = (1 - \alpha) \cdot V(\theta_{perceived}, \sigma_{motor}) + \alpha/2\pi$$

The model 'Bayes_var' is similar to the model 'Bayes', except that to account for the possibility of exposure effect to sensory uncertainty, we allow sensory uncertainty $\sigma_{sensory}$ to vary for each of the following angles 0°, ±16°, ±32°, ±48°, ±64°, resulting in five $\sigma_{sensory}$ free parameters.

The model 'Bayes_varmin' is similar to the model 'Bayes', except that we allow $\sigma_{sensory}$ to vary only for the most presented angles ±32°, resulting in two $\sigma_{sensory}$ free parameters ($\sigma_{sensory}$ at ±32°, and $\sigma_{sensory}$ at all other angles).

#### Response strategy models

We wanted to control for the possibility that the task behaviour might be explained by simple behavioural strategies that do not involve Bayesian integration. This class of models assumed that participants did not combine their expectations with sensory information, but relied on either of them alone on any given trial.

The first model, 'ADD1', assumed that estimations derived from prior expectations were simply sampled from a learnt prior distribution, $p_{expected}(\theta)$, which was parameterized as in Eq (4). However, on trials when participants did perceive motion direction, it was based solely on the sensory input, $p_{sensory}(\theta_{sensory}|\theta_{actual}) = V(\theta_{actual}, \sigma_{sensory})$.

Putting together the estimations derived from sensory input and the ones derived from learnt expectations, and the possibility of random estimations, the average distribution of estimation responses for a single participant is:

$$p(\theta_{estimate}|\theta_{actual}) = (1-\alpha)\cdot [(1-a(\theta))\cdot p_{sensory}(\theta_{sensory}|\theta_{actual}) + a(\theta)\cdot p_{expected}(\theta_{estimate})] * V(0,\sigma_{motor}) + \alpha, \quad (9)$$

where $a(\theta)$ determines the proportion of trials in which participants sampled from the acquired prior, $p_{expected}(\theta)$; asterisk ($*$) denotes convolution. The resulting 'ADD1' model has 9 free parameters ($\theta_{expected}$, $\sigma_{expected}$, $a(\theta)$ (which can take a different value for each of the 5 angles: 0°, ±16°, ±32°, ±48°, ±64°), $\sigma_{sensory}$ and $\alpha$).

The second model, 'ADD2', was the same as 'ADD1' except that it had more complex strategy for trials when participants relied on the prior: instead of sampling from the complete acquired prior distribution ranging from −180° to +180° (Eq. (4)), they sampled from only one half of it, negative (−180° to 0°) or positive (0° to +180°), depending on which side of the distribution the actual stimulus occurred on:

$$p_{expectedN}(\theta) = V(-\theta_{expected}, \sigma_{expected}) \quad (10)$$

$$p_{expectedP}(\theta) = V(\theta_{expected}, \sigma_{expected}) \quad (11)$$

Incorporating this into the distribution of estimation responses results in:

$$p(\theta_{estimate}|\theta_{actual}) = (1-\alpha)\cdot[((1-a(\theta)-b(\theta))\cdot p_{sensory}(\theta_{senssory}|\theta_{actual}) + a(\theta)\cdot p_{expectedN}(\theta_{estimate}) + b(\theta)\cdot p_{expectedP}(\theta_{estimate})] * V(0,\sigma_{motor}) + \alpha, \quad (12)$$

where asterisk ($*$) denotes convolution; $a(\theta)$ and $b(\theta)$ determine the proportion of trials in which participants sample from either negative or positive parts of the prior distribution, respectively.

Finally, we also considered two variations of the 'ADD1' and 'ADD2' models. These were identical to 'ADD1' and 'ADD2' except from setting $\sigma_{expected}$ to zero (i.e. no uncertainty); that is, on trials when perceptual estimates were derived only from expectations, they were equal to the mode of the learnt distribution. These models are referred to as 'ADD1_m' and 'ADD2_m'.

**Parameter estimation**

We used the performance in trials with the highest contrast level to estimate motor noise, $\sigma_{motor}$, for each individual. We assumed that at this level sensory uncertainty was close to zero ($\sigma_{sensory} \approx 0$). The motor noise was determined by fitting estimation responses at the highest contrast level to the distribution in Eq. (2) using the actual motion direction, $\theta_{actual}$, as the mean. The estimated motor noise was used in all the subsequent model fitting as a fixed parameter.

The rest of the free parameters of each model were estimated by fitting the response data from the two staircased contrast levels (~200 trials per participant). For each model with a set of free parameters M, we computed the probability distribution $p(\theta_{estimate}|\theta_{actual}; M)$ of making an estimate $\theta_{estimate}$ given the actual stimulus direction $\theta_{actual}$. For the response strategy models, by definition, the $p(\theta_{estimate}|\theta_{actual}; M)$ corresponds to average behaviour in the task (Equations 9 and 12). Bayesian models, on the other hand, explicitly model trial-to-trial variability in the posterior estimate, which in our case is the mean of the posterior (Eq. (6)). To relate this to the behavioural data we built a distribution of 1,000 samples for each presented angle (where each sample is the mean of the posterior obtained via Eq. (6) and perturbed by motor noise via Eq. (7) or (8)).

The parameters were estimated by maximizing the fit of the log likelihood function for the experimental data for each participant individually:

$$M = \text{argmax}_M \left[\sum_i^n \log\left(\text{p}(\theta_{estimate} = \theta_{i,data}|\theta_i)\right)\right], \quad (13)$$

where $\theta_{i,data}$ is participant's estimation response, $\theta_i$ is the actual presented motion direction on the ith trial and n is the number of trials. The maximum likelihood was found using *fminsearchbnd* function in Matlab, by minimizing negative log-likelihood. Parameters $\alpha$, $a(\theta)$ and $b(\theta)$ were bounded between 0 and 1, while $\theta_{expected}$,

$\sigma_{expected}$ and $\sigma_{sensory}$ were bounded from 0 to ∞. To reduce the possibility of convergence at a local maxima we constructed a grid of initial $\sigma_{expected}$ and $\sigma_{sensory}$ parameter values covering the range as found in previous studies ($\sigma_{expected}$ from 10° to 25° in 3° increments and $\sigma_{sensory}$ from 5° to 15° in 3° increments, which resulted in 22 different initializations). A set of parameters with the largest log-likelihood was selected as the best fit.

**Model Comparison**

To compare the model fits we used Bayesian Information Criterion (BIC), which approximates the log of model evidence (e.g., see Burnham and Anderson, 2004):

$$-2 \cdot \log(P(D|M)) \approx BIC = -2 \cdot \log(P(D|M, \Theta)) + k \cdot \log(n) \quad , \qquad (14)$$

where M is model, D is observed data and P (D|M, Θ) is the likelihood of generating the experimental data given the most likely set of parameters, Θ; k is the number of model parameters and n is the number of data points (or equivalently, the number of trials). BIC evaluates the model by how it fits the data by also penalizing for the number of parameters (i.e. model complexity) to avoid over-fitting. Lower BIC score indicates a better model.

**Parameter recovery**

To test the reliability of the parameter estimates of our winning model we performed parameter recovery. This allowed us to simultaneously test whether parameters are identifiable (e.g., whether likelihood and prior uncertainty is not correlated and can be distinguished) and whether having ~200 trials (the amount of low contrast trials in our data) for data fitting and using maximum likelihood estimation are sufficient to give reliable results.

First, we generated 100 sets of parameters (i.e. 100 synthetic individuals) by randomly sampling each parameter from a Gaussian distribution which had a mean and variance as the parameter estimates from the collected participant data. Second, for each set of parameters we simulated data for 200 trials with the winning model by randomly sampling from the estimation probability distribution, which, as for the behavioural data, was built from a 1000 posterior means (Eq. (6)), each perturbed by motor noise (Eq. (8)) Finally, we fitted the winning model to the simulated data. To evaluate the goodness of recovered parameters we computed the coefficient of determination ($R^2$) for a linear regression, which quantified how well the actual parameters predicted the recovered ones.

### Power calculations

**Power calculations (a-priori): Detecting effect of the prior on behaviour:**

We performed power calculations using an independent dataset from a previously published study that used the exact same task design but had 2 sessions of 850 trials (Chalk *et al.*, 2010). We used the first 567 trials of the first session for the power analysis, to conform to the same trial structure present in the current study.
'Bias' and 'Detection rate' were selected as the behavioural measures relevant for determining the effect of the prior on behaviour. Given these parameters, power calculations for 'Bias' and 'Detection rate' revealed that 18-20 subjects are required respectively to detect an effect with 80% power.

**Power calculations (a-priori): Detecting group differences:**

To our knowledge, the current task design has never been tested in a chronic Schizophrenia sample. We therefore turned to the literature to estimate the effect size between patients and controls in motion perception (Chen, 2011) and illusions (Dima et al., 2009; Dima et al., 2011).

We determined that 20 participants in each group should allow us to detect an effect size of at least 0.9 with 80% power, which is a smaller effect than previous effect sizes reported in motion perception (Chen et al., 1999a; Chen et al., 1999b), perceptual illusions (Dima et al., 2009; Dima et al., 2011), or cognition generally (Fioravanti *et al.,* 2012).
Based on this analysis we aimed to recruit approximately 25 subjects per group, which would permit to reliably detect the effect of: the prior on perception, as well as

differences in motion perception or illusions between groups, even accounting for a potentially large (e.g. 20%) exclusion rate based on task performance.
